# Active gaze behavior organizes V1 activity in freely-moving marmosets

**DOI:** 10.64898/2026.02.05.704079

**Authors:** Jingwen Li, Vikram Pal Singh, Alexander C. Huk, Jude F. Mitchell, Cory T. Miller

## Abstract

Human and nonhuman primates rely heavily on vision to actively explore and navigate their environment. Although primate visual cortex has been studied extensively in head-fixed animals, little is known about how the primate visual system supports natural, active vision in freely moving animals. Here, we address this gap in the primary visual cortex (V1) by leveraging a head-mounted eye-tracking system while simultaneously recording the activity of ensembles of single V1 neurons in freely moving marmosets. Our results reveal that primate neural activity is tightly driven by visual input and organized by the temporal structure of natural gaze behavior, and these gaze-related responses are largely abolished in the absence of visual input. We further show that distinct phases of gaze movement, i.e. rapid redirection (gaze shift) and subsequent stabilization (fixation), engage separable suppression and enhancement of the V1 responses. The enhancement during fixation was clearly linked to visual input. Moreover, the amplitudes of naturally-occurring gaze shifts organize RF input across successive fixations, and adaptively shapes fixation-associated enhancement. These findings define the dynamics in V1 that link natural gaze behavior and stimulus-driven responses in freely moving primates. The work opens a previously inaccessible but fundamental regime of primate vision and establishes freely moving paradigms as a foundation for understanding real-world visual processing during ethologically relevant behaviors.

**Significance Statement:** Vision in everyday life unfolds through active exploration that continuously reshapes the visual input reaching the brain. Yet nearly all we know about primate visual cortex derives from head-fixed paradigms that constrain these natural dynamics. Here, we present the first study of neuronal activity in primate primary visual cortex (V1) during fully unconstrained, freely moving visual exploration. We show that natural gaze behavior dynamically structures visual input and that V1 activity remains tightly anchored to visual signals during active exploration. By establishing freely moving primate paradigms as essential for understanding how vision operates in real-world conditions, this work provides a critical foundation for a more complete account of visual processing in the human and nonhuman primate brain.

## Introduction

Vision is the dominant sensory modality through which human and nonhuman primates perceive the world and guide behavior. A defining feature of vision is active sampling: rapidly scanning the environment to selectively acquire information through self-generated gaze shifts^1–6^. Despite increasing efforts to study visual cortex using natural scenes and free viewing paradigms^7–12^, visual cortical processing has not been examined under truly natural conditions in which primates are freely moving and actively exploring their environment. In such contexts, coordinated eye, head, and body movements generate rich, goal-directed sequences of visual behaviors that create rapid temporal dynamics on visual input that differ fundamentally from those produced in head-fixed paradigms^13–17^. These dynamics present unique challenges to the primate visual system and likely engage visual computations that are obscured under constrained conditions. This process is supported by the high-acuity fovea and continual redirection of gaze to behaviorally relevant locations uniquely in primates^18,19^. As a result, natural visual exploration produces meaningful statistics and dynamics of retinal input that are shaped by the primate’s own actions, profoundly impacting visual processing and perception. However, these computations remain largely unexplored due to the lack of experimental paradigms and datasets that examine visual processing in freely moving primates, as well as analytical frameworks capable of handling continuous, behavior-driven visual streams rather than trial-based, strictly controlled stimuli.

The rich patterns of eye, head, and body movements during natural exploration may also introduce non-visual signals from motor planning and corollary discharge that drive unique temporal and spatial statistics of visual input. Numerous studies have shown prominent movement-related modulation in rodent V1^5,20–27^, and head-fixed studies in primates suggest some role for eye-movement related modulation^28–31^. However, recent evidence suggests that, in contrast to rodents, movement-related influences on primate V1 are minimal under constrained conditions^32,33^. And while rich neural dynamics have been observed with gaze shifts in freely moving mice^5^, they have not been fully examined when primates are freely moving. Whether fully unconstrained behavior could elicit more prevalent or stronger non-visual modulation in primate visual cortex remains an open question.

To address these questions, we established a paradigm for studying primate vision in completely unconstrained conditions and examined visual processing in V1 as marmoset monkeys freely explored their environment. We found that V1 neurons respond to gaze shifts with a suppression followed by an enhancement, consistent with previous results in mice and primates^5,34,35^. However, these responses depended strongly on visual input and showed minimal contribution from non-visual signals, in striking contrast to mouse V1. We further demonstrated that suppression and enhancement correspond to distinct phases of continual gaze behavior: suppression accompanies the rapid retinal changes during gaze shifts, whereas enhancement encodes stimuli within the neuron’s receptive field (RF) during fixations when retinal input is stabilized. Moreover, we demonstrated that the amplitude of natural gaze shifts organizes RF stimuli across successive fixations, such that larger gaze shifts produce more distinct visual input and stronger fixation-associated enhancement. Together, these findings reveal how primate V1 processes visual information during fully unconstrained, real-world contexts, which is tightly anchored to visual signals and the temporal structure of natural gaze dynamics.

## Results

### V1 responses are structured by gaze during active exploration

We recorded the activity of ensembles of single neurons in V1 of marmoset monkeys freely moving in a visually enriched arena, while simultaneously tracking the eye, head, body, and visual scene (Fig. 1a; Table S1; see Methods). During active exploration, the monkeys visually scanned the environment through frequent alteration between redirecting and stabilizing gaze toward points of interest (Fig. S1a). Such gaze shifts and fixations were achieved through coordinated eye and head movements: eye and head movements were in the same direction during gaze shifts, whereas they were negatively correlated during fixations due to the vestibulo-ocular reflex (Fig. 1b).

**Figure 1.**
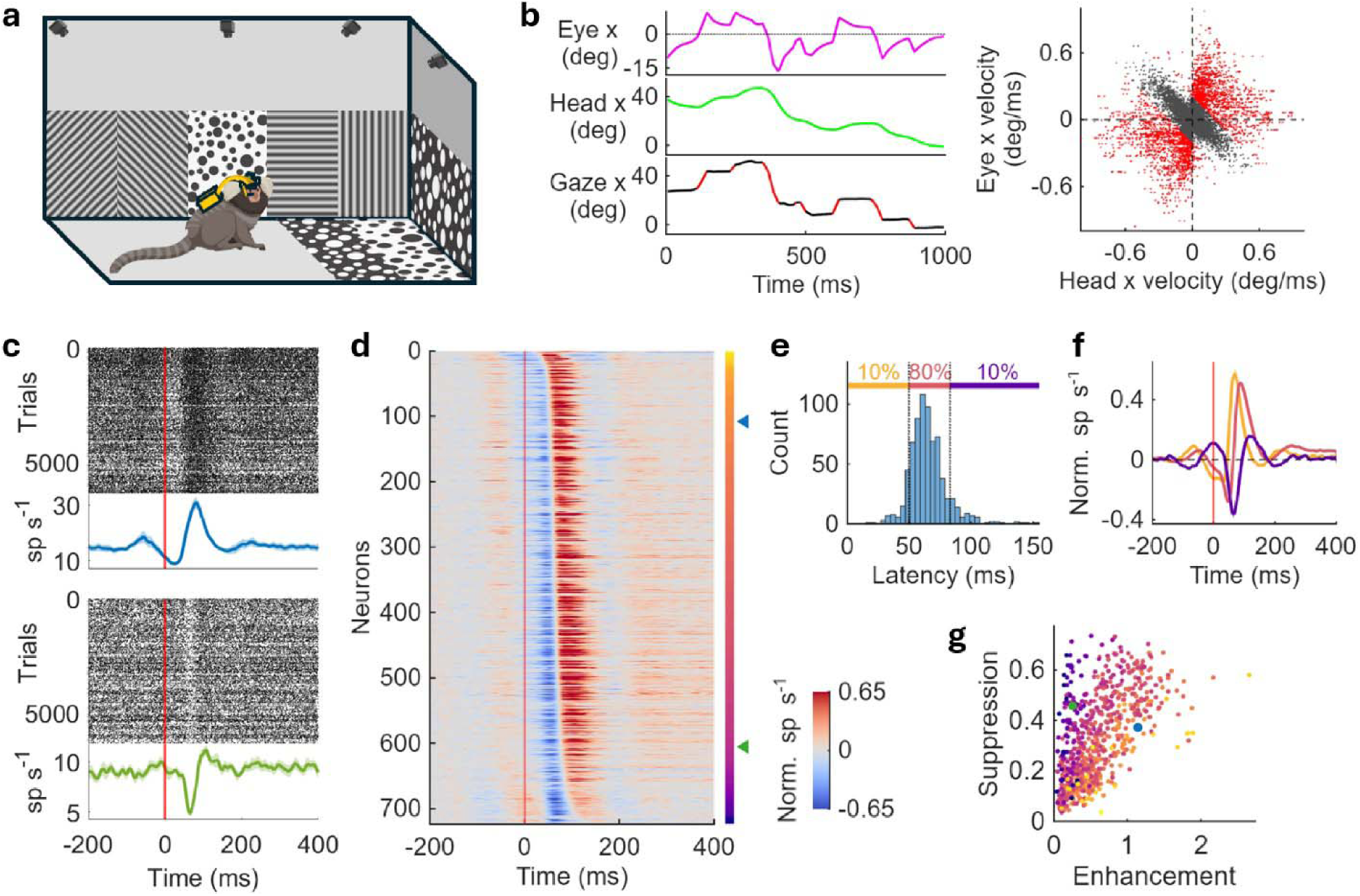
V1 neurons robustly respond to gaze shifts with a suppression followed by enhancement in a temporal variability of latencies. **a)** Experimental paradigm. The monkey freely moves in an arena decorated with dots and gratings. **b)** Eye and head coordination. Left: horizontal eye (top), head (middle) and gaze (bottom) position. Right: horizontal eye velocity versus horizontal head velocity. Eye and head velocities are negatively correlated during gaze fixation (black); the negative correlation is broken during gaze shifts (red). **c)** Two example neurons’ raster plots and PSTHs aligned with gaze-shift onset. Both neurons exhibit a suppression followed by enhancement. Dots represent spikes in raster plots; lines and shades represent mean and standard error in PSTHs, respectively. Red lines indicate gaze-shift onsets. **d)** Normalized PSTHs aligned with gaze-shift onset in the V1 population sorted by response latencies. The population shows a robust response in a temporal variability. The vertical bar on the right indicates latencies from early (yellow) to late (purple). The two example neurons in **c** are indicated by the blue and green triangle. **e)** Histogram of response latencies. Most neurons fall in the middle range of latencies. 0.1 and 0.9 quantities are indicated by dotted lines and labeled in different colors above the histogram. **f)** Population-average normalized PSTHs of neurons grouped by 0.1 and 0.9 quantities of latencies as in **e** with the same color. Enhancement is more prominent in earlier neurons and suppression is more prominent in later neurons. Shades represent standard error. **g)** Suppression amplitude versus enhancement amplitude. Each dot represents a neuron with color indicating latency as in **d**. The suppression/enhancement ratio is larger in later neurons than earlier neurons. The two example neurons in **c** are indicated by the blue and green dot.

We first examined how V1 neurons respond to gaze shifts. Raster plots and peristimulus time histograms (PSTHs) of two example neurons showed a suppression followed by an enhancement in firing rate after gaze-shift onset (Fig. 1c). The two neurons differed in the amplitudes, durations, and latencies of their suppression and enhancement responses. Across the 743 neurons we recorded from two monkeys, 71.3% (n=530) and 86.3% (n=641) exhibit significant suppression and enhancement, respectively (amplitudes exceeding ±3 SE of baseline; see Method). A heatmap of normalized PSTHs aligned with zero-crossing point between suppression and enhancement following gaze-shift onsets revealed a temporal variability of response latencies across the population (Fig. 1d), with most neurons falling within the middle range of the latencies (Fig. 1e). Sub-population averages of normalized PSTHs showed that early-latency neurons displayed stronger enhancement, whereas late-latency neurons exhibited more prominent suppression (Fig. 1f). Consistently, suppression and enhancement amplitudes varied systematically across neurons, with later-responding neurons exhibiting a larger suppression-to-enhancement ratio (Fig. 1g). These findings were consistent across animals (Fig. S1b-d) and with previous studies examining only eye movements in head-fixed primates^5,30,31^. Together, these results demonstrate that natural gaze shifts resulting from both eye and head movements evoke a robust, time-structured response in marmoset V1, characterized by a suppression followed by an enhancement that unfolds in a temporal variability across the population.

### Gaze-shift initiated responses in V1 depend on visual input

To determine whether V1 responses to gaze shifts depended on visual input or non-visual signals, we performed the experiment in complete darkness, eliminating visual input while gaze shifts persisted (Fig. 2a, S2a-c). Strikingly, both suppression and enhancement were largely abolished in darkness (Fig. 2b). Population-averaged normalized PSTHs showed a dramatic reduction in response amplitude relative to the light condition (Fig. 2c). Although weak residual responses were observed in subpopulation averages, their amplitudes were substantially smaller than in the light (Fig. 2d). Only 6.3% (n=19) and 9.9% (n=30) out of 304 neurons exhibited significant suppression and enhancement in darkness, respectively (±3 SE criterion, no correction for multiple comparisons; see Methods; examples shown in Fig. S2d). Correspondingly, the systematic relationship between suppression and enhancement amplitudes associated with response latencies was disrupted in darkness (Fig. 2e, S2e). Although firing rates were generally lower in darkness, the absence of gaze-shift responses was not due to a lack of spiking activity, as most neurons maintained appreciable firing rates in the dark (Fig. 2f). Sorting neurons by their response latencies in the dark did not preserve the temporal structure of responses observed in the light, indicating that the weak residual responses in the dark do not systematically follow the response dynamics observed in the light (Fig. S2f–g). Together, these findings indicate that gaze-shift responses in marmoset V1 depend substantially on visual input, rather than being driven primarily by non-visual modulation. This sharply contrasts with V1 in freely moving mice where a widespread suppression persists in darkness^5^ and when retinal input is pharmacologically silenced^27^, suggesting fundamentally different contributions of non-visual signals to rodent versus primate V1 (see Discussion).

**Figure 2.**
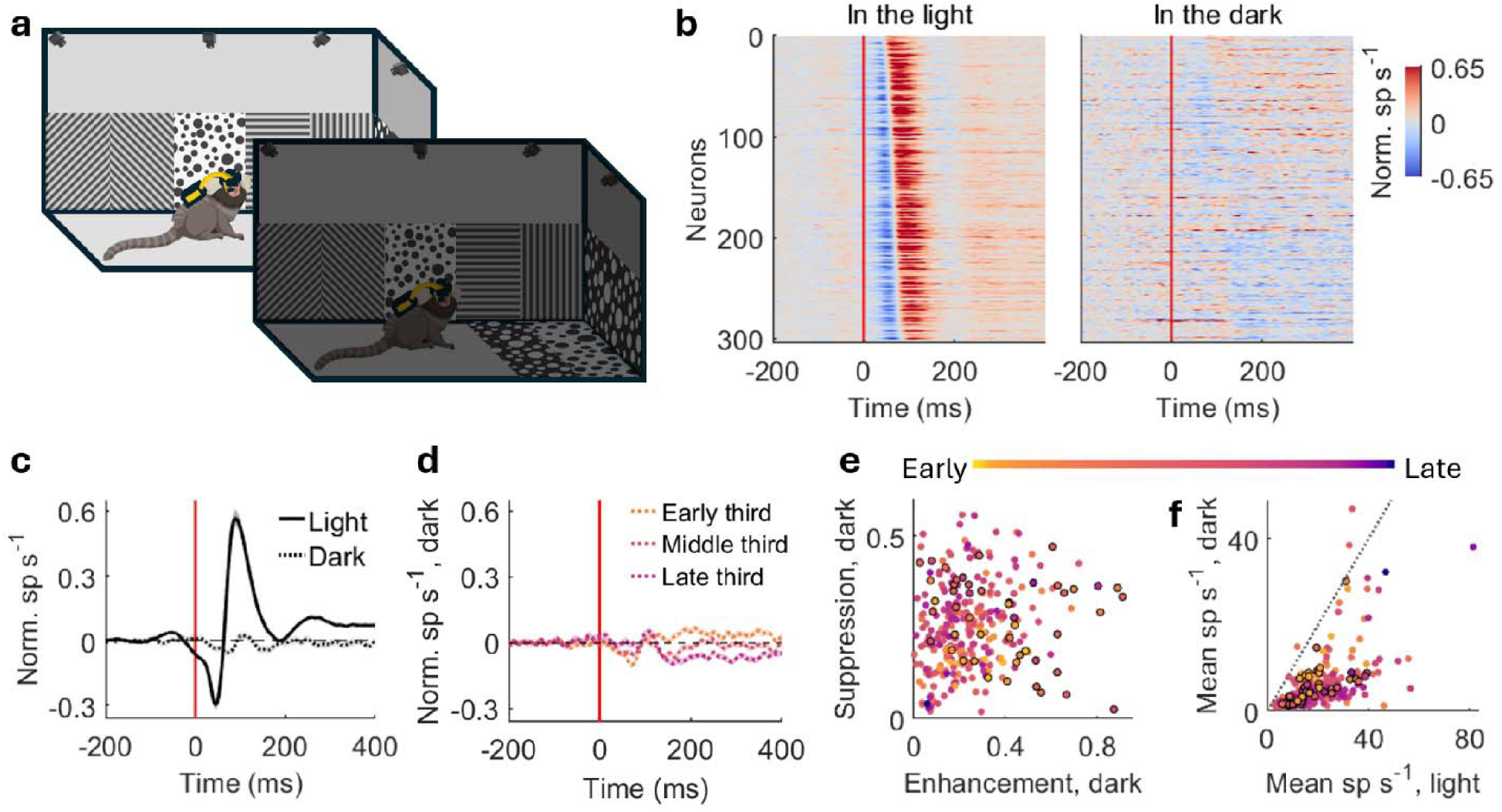
V1 responses to gaze shifts rely on visual inputs. **a)** Experimental paradigm. The monkey freely moves in the light and dark. **b)** Normalized PSTHs aligned with gaze-shift onset in the light (left) and dark (right) with the same order sorted by response latencies in the light. The suppression, enhancement, and temporal variability of latency in the light are vastly disrupted in the dark. Red lines indicate gaze-shift onsets (same in **c** and **d**). **c)** Population-average normalized PSTHs aligned with gaze-shift onset in the light (solid line) and dark (dotted line). The response to gaze shifts scarcely exists in the dark. Shades represent standard error. **d)** Population-average normalized PSTHs in the dark grouped by early, middle, and late third neurons as the same order in **b**. The little remaining suppression and enhancement in the dark are found in different groups of neurons. Shades represent standard error. **e)** Suppression amplitude versus enhancement amplitude in the dark. Each dot represents a neuron with color indicating latency as in Fig. 1d. Neurons with significant suppression or enhancement in the dark (see Method) are indicated in black circles. The pattern of larger suppression/enhancement ratio for later neurons in the light (Fig. 1g) is disrupted in the dark. **f)** Mean spike rate in the dark versus in the light. Each dot represents a neuron with color and black circle same as in **e**. The black dotted line represents equality. Most neurons maintained sufficient spiking activity in the dark.

### Distinct phases of gaze behavior elicit separable response components in V1 neurons

A key feature of active exploration in freely moving marmosets is the frequent and continuous alternation between gaze shifts and fixations, spanning a wide range of amplitudes and durations that occur with conjugate head+eye gaze shifts^17^. Visual processing must therefore accommodate considerable temporal variability. To elucidate the nature of visual responses elicited by the widely variable dynamics of gaze shifts during natural behavior, we re-examined raster plots and PSTHs aligned to gaze-shift onsets and fixation onsets separately (Fig. 3a). This analysis allowed us to dissociate response components associated with different phases of gaze behavior. Raster plots sorted by gaze-shift duration revealed that suppression was tightly time-locked to gaze-shift onset, whereas enhancement was time-locked to fixation onset. Consistent with this dissociation, enhancement amplitudes were larger in fixation aligned PSTHs than in gaze-shift aligned PSTHs. Across the population, enhancement amplitudes were consistently larger in fixation aligned than gaze-shift aligned PSTHs (Fig. 3b). Suppression amplitudes, in contrast, showed a smaller but significant bias for the opposite pattern, with stronger suppression when time-locked to saccade onset than fixation onset. Thus, the initial suppression near gaze shifts appears associated with the onset of the shift, while the later enhancement appears better associated with fixation after the shift.

**Figure 3.**
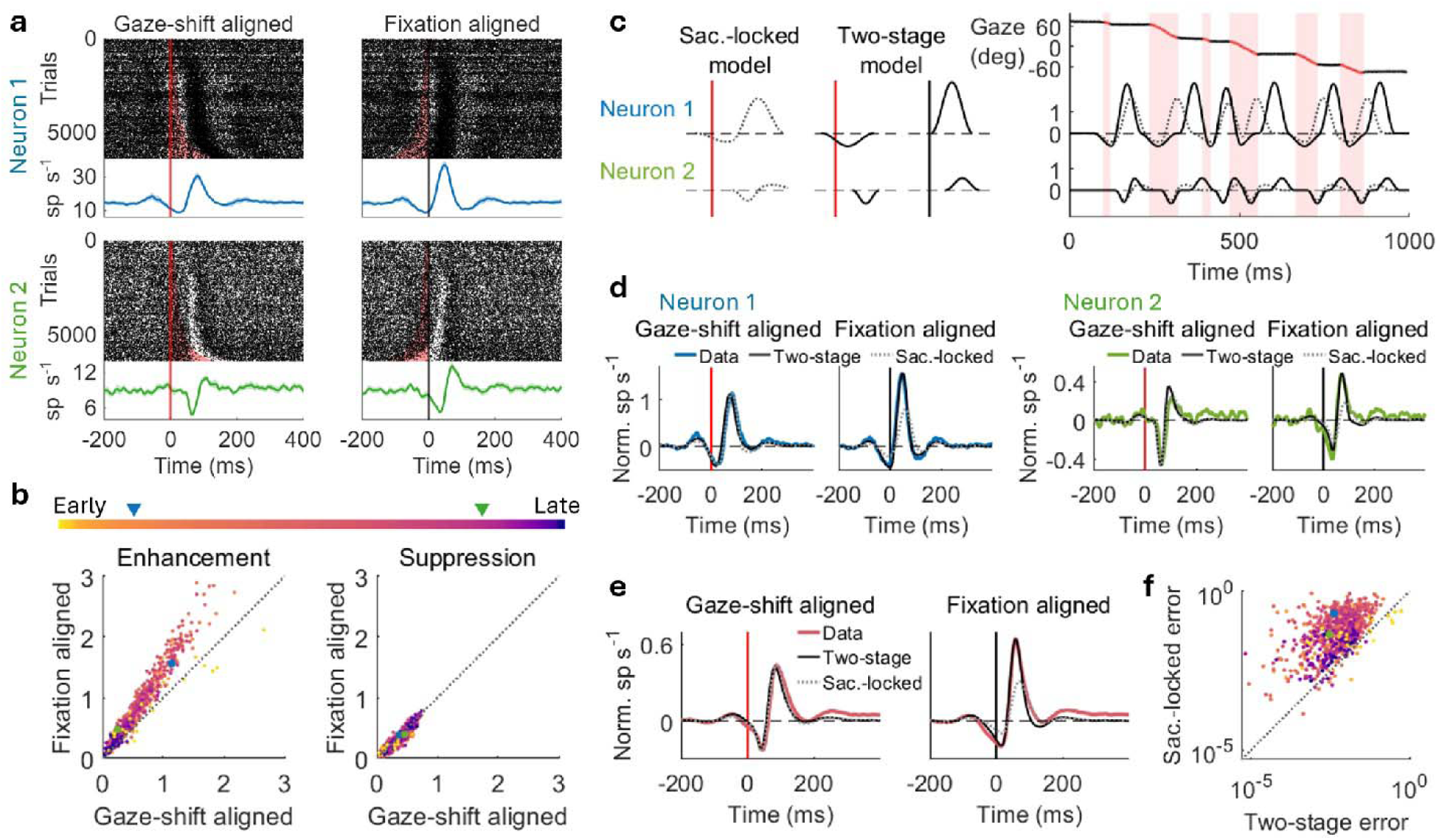
Suppression and enhancement are associated with gaze shift and fixation, respectively. **a)** Raster plots and PSTHs aligned with gaze-shift onsets (left) and fixation onsets (right) for the two example neurons in Fig. 1a. Trials are sorted by gaze-shift durations (red shaded area). Suppression is time-locked to gaze-shift onsets in raster plots and exhibits larger amplitudes in gaze-shift aligned PSTHs, whereas enhancement is time-locked to fixation onsets in raster plots and exhibits larger amplitudes in fixation aligned PSTHs. Dots represent spikes in raster plots; lines and shades represent mean and standard error in PSTHs, respectively. Red lines and black lines indicate gaze-shift and fixation onsets, respectively (same in **c**, **d** and **e**). **b)** Fixation aligned versus gaze-shift aligned enhancement amplitude (left) and suppression amplitude (right). Each dot represents a neuron with color indicating latency as in Fig. 1d. The two example neurons in **a** are indicated by blue and green dots and triangles. **c)** Parametric models describing neural responses to gaze movements (see Method). Saccade-locked model: suppression is followed by enhancement after gaze shifts (gray dotted line). Two-stage model: suppression and enhancement are associated with gaze shifts and fixations, respectively (black solid lines). The two models are applied on each gaze shift and fixation continuously throughout the recording for the two example neurons. Red shaded areas indicate gaze shifts. As gaze-shift durations vary, the saccade-locked model and two-stage model exhibit different neural responses to gaze movements. **d)** Normalized PSTHs aligned with gaze-shift onset and fixation onset obtained from the best-fit saccade-locked model (gray dotted lines) and two-stage model (black solid lines) compared to data (colored lines) for the two example neurons. The two-stage model outperforms the saccade-locked model in fixation aligned PSTHs. **e)** Population-average normalized PSTHs aligned with gaze-shift onset (left) and fixation onset (right) obtained from the best-fit saccade-locked model (gray dotted lines) and two-stage model (black solid lines) compared to data (coral lines). **f)** Error of the saccade-locked model versus the two-stage model on log–log axes (see Method). Each dot represents a neuron with the same color in **b**. The two-stage model outperforms saccade-locked model for most neurons. The two example neurons in **a** are indicated by blue and green dots.

We formally tested this dissociation using two parametric models of gaze-shift responses (Fig. 3c; see Methods). In the “saccade-locked” model, both suppression and enhancement were linked to gaze-shift onsets. In the “two-stage” model, suppression was linked to gaze-shift onset, and enhancement to subsequent fixation onset. When applied to the continuous sequence of natural gaze behavior, the models yielded distinct neural response predictions, especially when gaze-shift durations varied considerably for larger amplitude shifts (Fig. 3c). The wide range of gaze-shift sizes in natural vision yield a broad span of latencies between shift onset and subsequent fixation, and thus allows us to dissociate shift- and fixation-related response components. By contrast, head-fixed saccades fall within a limited range of amplitudes where shift duration is less variable^5,31^. We next derived best-fit PSTHs from each model’s continuous response predictions and compared them to the data-obtained PSTHs. While both models recapitulated gaze-shift aligned PSTHs reasonably well in the two exemplar neurons, only the two-stage model successfully captured the larger enhancement observed in fixation aligned PSTHs (Fig. 3d). These exemplars were consistent with effect at the population level (Fig. 3e). Model performance quantified using mean-square error of suppression and enhancement amplitudes in gaze-shift and fixation aligned PSTHs showed that the two-stage model outperformed the saccade-locked model for the vast majority of neurons (Fig. 3f). These results validated that suppression and enhancement are two temporally distinct components linked to gaze shifts and fixations, respectively, rather than a single saccade-locked response. While the saccade-locked model has sufficiently described results from head-fixed monkeys in previous studies^31^, the two-stage mechanism emerges clearly in the continuous, unconstrained gaze behavior of freely moving marmosets.

### Fixation-associated enhancement reflects visual processing of retina input in V1

Having characterized how dynamic gaze behavior structures V1 responses temporally, we next asked how visual processing is integrated into these responses. Given that enhancement is fixation-associated, we hypothesized that this enhancement reflects the processing of visual input during fixations. To test this hypothesis, we first mapped each neuron’s receptive field (RF) and tuning in a head-fixed paradigm where monkeys were presented flashing dots and drifting gratings (Fig. 4a). For each neuron, we identified its RF location and computed a weighted-mean preferred input for stimulus orientation and spatial frequency using the frequency domain magnitude spectrum derived from its tuning^36^ (Fig. 4b; see Methods). The monkeys were then allowed to freely explore the arena while the same neurons were recorded. During active exploration, the visual input falling within the neuron’s RF was extracted from the world-view camera and converted into a frequency-domain representation for each fixation (Fig. 4c). We directly compared each neuron’s preferred RF input while head-fixed with the current RF input while freely-moving to obtain a predicted RF drive across gaze fixations (Fig. 4d).

**Figure 4.**
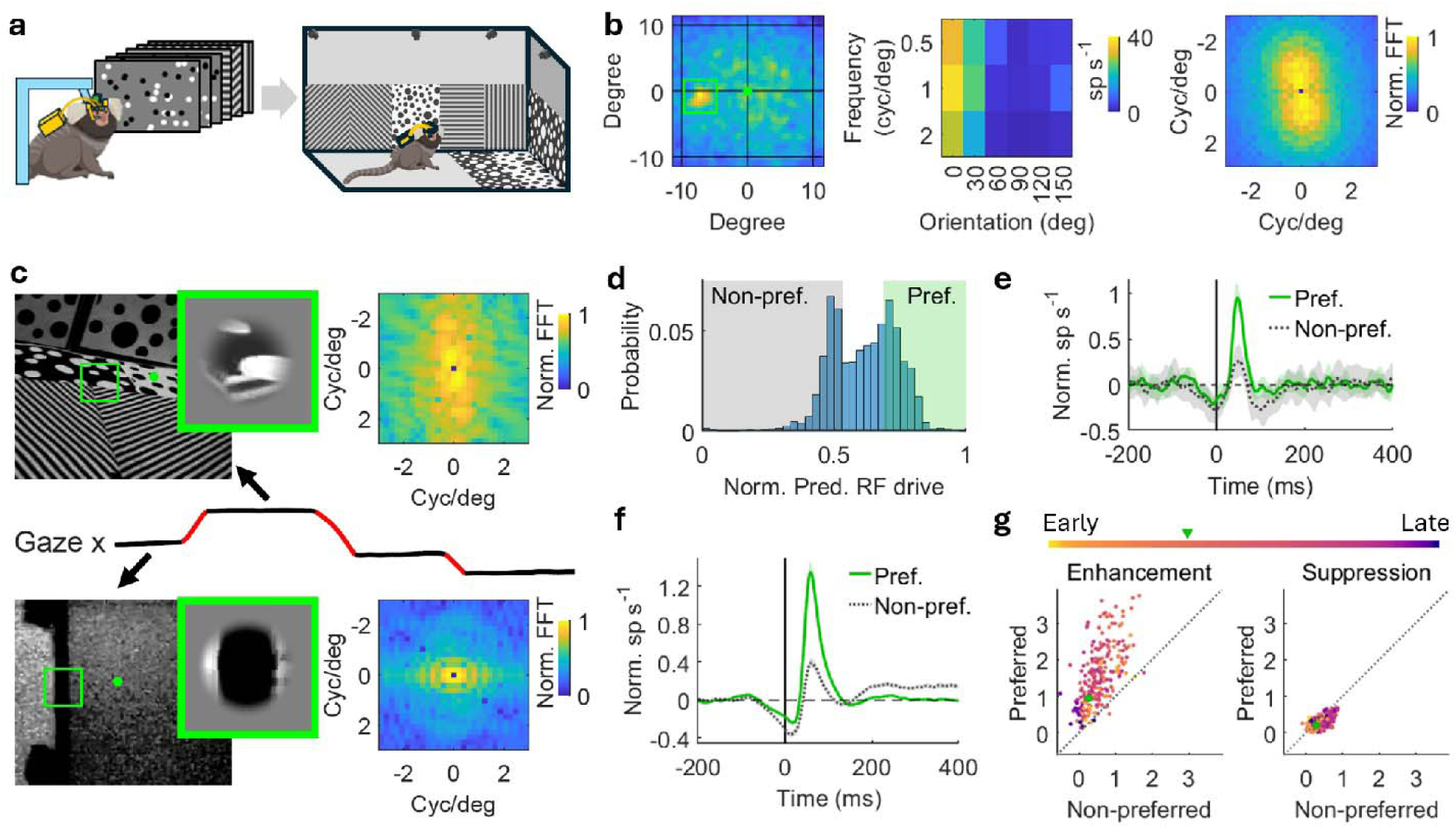
Enhancement associated with fixation is driven by the alignment of visual inputs during fixation to the V1 neuron’s preferred tuning. **a)** Experimental paradigm. The monkey is head-fixed and shown flashing dots and drifting gratings before being brought to freely move in the arena. **b)** Receptive field (left), tuning property (middle), and weighed mean normalized FFT log-magnitude spectrum of an example neuron’s tuning (right). **c)** Visual inputs of two example fixations for the example neuron in **b**. Images are captured by the world-view camera on the head-mounted eye tracker (left). The green dot and box in the image represent the gaze fixation point and receptive field of the neuron, respectively. The visual input in the receptive field is cropped out and added a raised cosine filter (middle). FFT is applied to the visual input (right) and a predicted RF drive is obtained by comparing the FFT log-magnitude spectrum of the neuron’s preferred RF input with the current RF input (see Method). The example at the top has a larger predicted RF drive than the example at the bottom. **d)** Distribution of the normalized predicted RF drive of visual input across fixations. Fixations with a predicted RF drive >2/3 quantile (gray shaded area) and <1/3 quantile (green shaded area) are selected in the preferred and non-preferred group, respectively. **e)** Normalized PSTHs aligned with fixation onsets grouped in preferred (solid line) and non-preferred (dotted line) visual inputs for the example neuron in **b**. Shades represent standard error. The black line indicates fixation onsets (same in **f**). **f)** Population-average normalized PSTHs of the V1 population aligned with fixation onset of preferred (solid line) and non-preferred (dotted line) visual inputs. The difference of enhancement is larger than the difference of suppression. Shades represent standard error. **g)** Enhancement (left) and suppression (right) amplitude of preferred visual-input PSTHs versus non-preferred visual-input PSTHs. Each dot represents a neuron with the same color in **f**. The example neuron used in **b**-**e** is indicated by the green dot.

Fixations were labeled as preferred (top tertile) or non-preferred (bottom tertile) based on their predicted RF drive, and fixation-aligned neural responses were examined under these two conditions. As shown in an example neuron, enhancement was substantially larger for preferred versus non-preferred inputs in fixation aligned PSTHs, whereas suppression remained unchanged (Fig. 4e). Population-averaged PSTHs showed the same pattern: preferred RF inputs evoked much stronger enhancement with only modest effects on suppression (Fig. 4f).

Enhancement amplitudes were consistently higher for preferred compared to non-preferred inputs across neurons, whereas suppression amplitudes showed only a slight bias toward non-preferred inputs (Fig. 4g). These results indicate that enhancement reflects visual processing of the RF input during fixation, specifically how well the current visual input matches the neuron’s preferred tuning. In contrast, suppression is largely independent of the visual content but rather reflects the visual transient generated by the gaze movement, and potentially, nonvisual modulation.

### Naturally-occuring gaze-shift amplitudes sculpt the interaction of receptive field input and fixation-associated enhancement

Rather than being restricted to discrete and well-defined preferred and non-preferred stimuli, RF input varied continuously (and in many dimensions) across fixations during natural visual exploration in freely moving marmosets. Therefore, following the previous analysis for RF input, we further characterized fixation-associated enhancement as a function of the predicted RF drive (Fig. 5a-c; see Methods). Across all neurons and recording sessions, normalized predicted RF drive spanned a broad continuum (Fig. 5a, bottom). Along this continuum, lowest predicted RF drive values primarily reflected low-contrast RF inputs, whereas the highest values were dominated by high-contrast inputs. Within the intermediate range, predicted RF drive represented the degree of alignment between the spatial structure of the RF visual input and the neuron’s tuning (Fig. 5a, top). We therefore restricted subsequent analyses to the 10th–90th percentile of predicted RF drive values. For an example neuron, firing rates during fixation-associated enhancement increased with predicted RF drive (Fig. 5b). This relationship was also observed at the population level, where 84% of neurons (211 out of 251) exhibited a significant positive linear relationship between predicted RF drive and fixation-associated enhancement (Fig. 5c).

**Figure 5.**
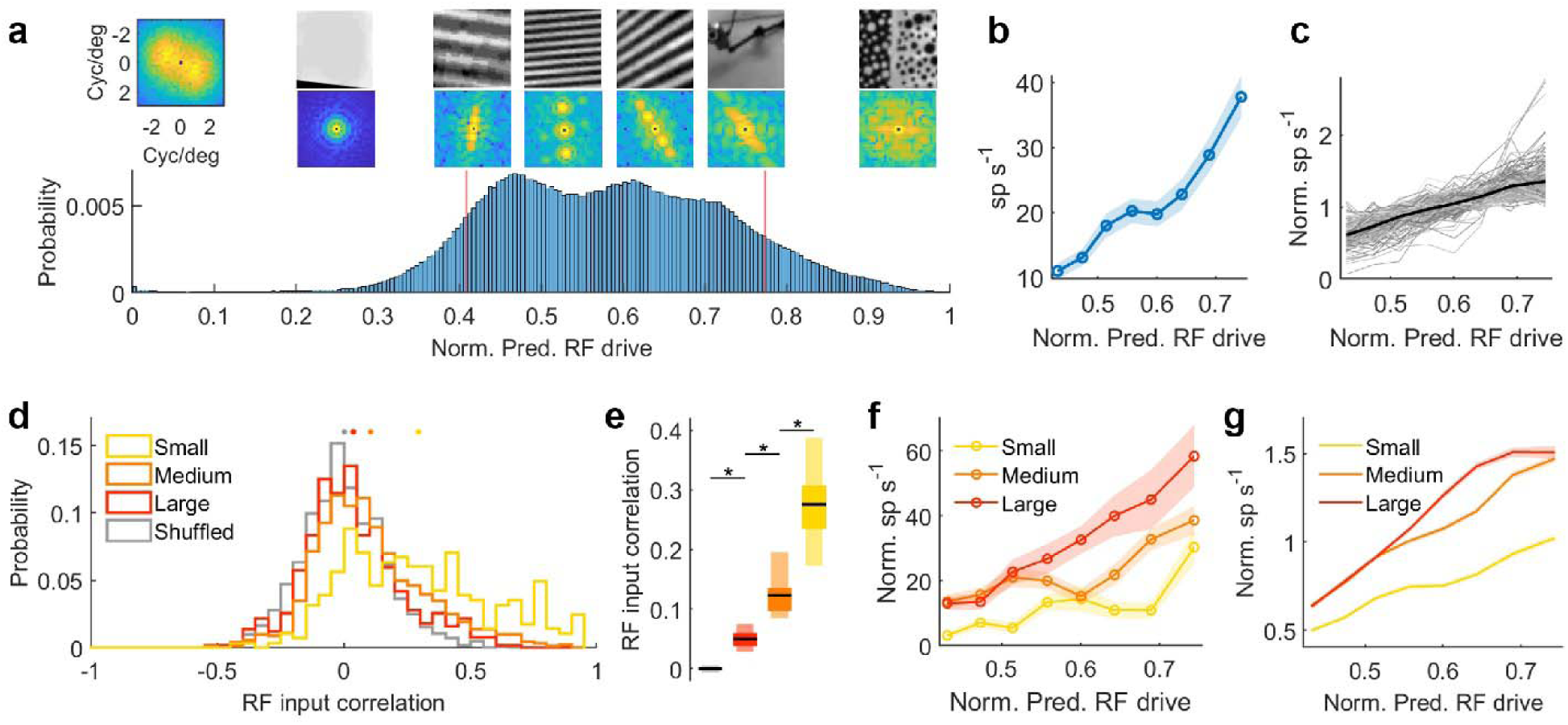
Gaze-shift amplitude organizes RF visual input across successive fixations and adaptively modulates fixation-associated enhancement in V1. **a)** Distribution of normalized predicted RF drive across fixations, pooled across all sessions and neurons. Upper left: tuning of an example neuron in the frequency domain. Upper right: example RF input stimuli and their corresponding frequency-domain representations at representative predicted RF drive values above the histogram. The red vertical lines indicate the predicted RF drive range used for the following analyses. Predicted RF drive values below the lower bound largely correspond to low-contrast stimuli, whereas values above the upper bound are typically associated with high-contrast stimuli. Predicted RF drive values within the selected range primarily reflect the degree of alignment between the RF visual input and the neuron’s tuning. **b)** Mean firing rate at fixation-associated enhancement as a function of normalized predicted RF drive for the example neuron. Shade represents standard error. Mean firing rate increased with the normalized predicted RF drive. **c)** Normalized firing rate as a function of normalized predicted RF drive for neurons exhibiting a significant positive linear relationship (p<0.05). Gray lines show individual neurons; black line represents the population mean. Firing rates are normalized to each neuron’s mean response across normalized predicted RF drive bins. **d)** Marginal distributions of RF input correlation between consecutive fixations following small (yellow), medium (orange), and large (red) gaze shifts for an example session. The distribution of RF input correlation between shuffled fixations is shown for comparison (gray). Dots above show the mean of each distribution. Consecutive fixations following small gaze shifts exhibit higher RF input correlation than those following medium or large gaze shifts. **e)** Statistics of RF input correlation across recording sessions. Box plots show the mean (black line), 25th–75th percentile (wide box), and 10th–90th percentile (thin box) for each group (colors as in **d**). RF input correlation increases with decreasing gaze-shift amplitude. All pairwise comparisons were significant (paired t-test, p<0.05). **f)** Mean firing rate at fixation-associated enhancement as a function of normalized predicted RF drive for an example neuron, grouped by the amplitude of the preceding gaze shift. Curves correspond to small (yellow), medium (orange), and large (red) gaze shifts. Shades represent standard error. Across all predicted RF drive values, firing rates increase with larger amplitude of the preceding gaze shift. **g)** Population average of normalized firing rate following fixation as a function of normalized predicted RF drive, grouped by the amplitude of the preceding gaze shift. Firing rates are normalized to each neuron’s mean response across preceding gaze-shift amplitudes and normalized predicted RF drive bins. Shades represent standard error. The population average exhibits the same pattern as the example neuron in **f**.

We next examined how natural gaze behavior structures the sequence of RF input across successive fixations and whether this organization influences visual processing. We found that the correlation between consecutive RF inputs systematically depended on the amplitude of the intervening gaze shift. In an example recording session, consecutive fixations following small gaze shifts exhibited substantially higher RF input correlation than those following medium or large gaze shifts (Fig. 5d). This relationship was consistent across recording sessions, with RF input correlation decreasing as gaze-shift amplitude increased (Fig. 5e). We therefore hypothesized that fixation responses would depend not only on the current RF input but also on the correlation between consecutive RF inputs established by the preceding gaze shift.

Analyses showed that for a given level of predicted RF drive, firing rates during fixation-associated enhancement were higher following larger gaze shifts than smaller gaze shifts in the example neuron (Fig. 5f), thereby confirming with our prediction. The same pattern was observed at the population level (Fig. 5g, S4c), demonstrating that fixation-associated enhancement is jointly determined by the match between the current RF input and neuron’s tuning, together with the correlation between consecutive RF inputs shaped by the preceding gaze shift. These findings show that natural gaze behavior organizes both the spatial and temporal structure of retinal input, with gaze-shift amplitude structuring the correlation between visual input across successive fixations and adaptively modulating V1 responses during subsequent fixation.

## Discussion

In the present study, we examined how marmoset behavior influences the organization of visual information in the primary visual cortex during freely-moving exploration. We found that V1 neurons respond to gaze behavior with a temporally structured pattern consisting of suppression during gaze shifts followed by enhancement during subsequent fixations. Crucially, these responses depended strongly on visual input and were largely absent in darkness. This indicates that, in freely moving primates, V1 primarily encodes rapidly changing retinal input associated with natural gaze behavior rather than non-visual movement-related signals. Overall, our findings establish that primate V1 is organized by natural gaze behavior through a two-stage mechanism that dynamically structures visual information across space and time during freely moving exploration, with gaze-shift amplitude organizing receptive-field input across successive fixations and shaping fixation-associated enhancement.

The dense temporal structure of natural gaze behavior both shapes neural population responses and reveals mechanisms that are not evident under constrained conditions. First, the frequent and rapid succession of gaze shifts gives rise to weak suppression and enhancement components flanking the main response (Fig. 1d). These flanking features are recapitulated by the modeled PSTHs (Fig. S3c). Because the model incorporates only a single suppression and enhancement response while incorporating the actual sequence of gaze events observed during the recording, these flanking features arise naturally from the frequent alternation of gaze shifts and fixations rather than reflecting distinct neural mechanisms. Second, saccade-locked descriptions have been traditionally used to characterize saccadic suppression in head-fixed primates^31^. While the responses reported here are consistent with visual mechanisms previously described in head-fixed conditions, under these constrained conditions, the two-stage mechanism is largely obscured because saccade amplitudes and durations are limited by the range of eye movements. As a result, PSTHs aligned to gaze-shift onset and fixation onset appear nearly identical. In freely moving marmosets, however, gaze shifts span a much wider range of amplitudes and durations due to coordinated eye-head movements. This variability reveals a clear temporal dissociation between suppression during gaze shifts and enhancement during subsequent fixations captured by the two-stage model (Fig. S3d-f), which emerges as a salient feature of V1 processing during natural, real-world vision. Similar temporal dissociation has been reported in studies using simulated or controlled saccades, suggesting that these dynamics reflect an active reorganization of visual processing around eye movements^37–42^.

Third, we found that gaze-shift amplitude systematically determines the similarity of receptive-field input across successive fixations. Small gaze shifts tend to preserve visual input across fixations, whereas large gaze shifts introduce more distinct input. Correspondingly, fixation-associated enhancement is adaptively modulated by the amplitude of the preceding gaze shift, indicating that V1 responses are shaped not only by the current visual input but also by the recent sequence of visual input generated through natural gaze behavior.

Only a small subset of neurons with the earliest response latencies deviated from the suppression followed by enhancement pattern. Among this population, 6 of the total 633 neurons exhibited instead a direct enhancement that begins after gaze-shift onset (Fig. S3a-b), resulting in equal or even larger enhancement amplitudes in gaze-shift aligned compared with fixation aligned PSTHs (Fig. 3b). These neurons may be responding directly to the visual transient and the retinal motion induced from saccade onset itself, irrespective of any specific orientation or spatial frequency content in the following fixation epoch. If so, they would be expected to be segregated to layers of V1 sensitive to motion receiving magnocellular inputs sensitive to high temporal frequencies and motion (Layer 4B, 4Calpha), which would suggest our recording approach may have predominantly sampled rather from layers driven by parvocellular inputs (layers 2/3, 4Cbeta), which constitute the majority of primate V1. One limitation of the current study is that we do not have information about the laminar location of the recorded neurons. Therefore, testing whether suppression, enhancement, and their tuning properties differ across cortical layers will require future recordings with laminar probes that allow precise estimation of neuronal depth.

Beyond temporal structure, an important implication of our findings concerns how experimental constraints shape gaze-related modulation in primate V1. Previous studies in head-fixed macaques reported modulation of V1 activity by gaze direction^43,44^. However, in head-fixed paradigms, eye position is decoupled from head movements, such that gaze direction depends solely on eye position. This constraint fundamentally differs from gaze behavior in freely moving primates, where coordinated eye-head movements dynamically redirect gaze and stabilize fixation near the center of the oculomotor range^17^. As a result, gaze direction in natural behavior reflects an integrated eye-head strategy rather than eye position alone, rendering these prior observations not directly comparable to our results. This distinction may also help reconcile differences in the contribution of non-visual signals to V1 activity. In head-fixed macaques, residual saccade-related responses in darkness have been reported at the population level in V1^31^. In contrast, we observed that gaze-related responses in freely moving marmoset V1 were largely abolished in darkness. One possibility is that the residual responses observed under head-fixed conditions reflect eye-position related or other extraretinal signals that are amplified by decoupling of eye and head movements. However, the considerable head and body size differential between macaques and marmosets, and other species differences, may also contribute to the observed differences. Whether such signals arise from constrained conditions or reflect mechanisms that are suppressed or absent during natural eye-head coordination remains an important question for future work.

Our light–dark manipulation showing that V1 responses are largely abolished in darkness suggests that non-visual signals have little to no contributions at this early stage of the primate visual hierarchy. These findings stand in sharp contrast to findings in freely moving mice, where substantial non-visual modulation persists in V1 without visual input^5,27,45^. Consistent with our observations, recent studies showed that locomotion has only limited modulation on marmoset V1 activity^32^ and that spontaneous movements have minimal effects on macaque V1 responses^33^. While gaze-related non-visual signals appear to play a minimal role in primate V1, they may impose a stronger influence in higher-order primate visual areas. For instance, saccadic suppression has been consistently observed in darkness in primate area V4^46,47^, supporting the idea that extraretinal modulation may emerge more prominently at later stages of visual processing. These differences underscore distinct V1 coding strategies in freely moving primates compared to rodents that likely reflect broader circuit and computational mechanisms underlying vision^48,49^.

Together, our findings advance the understanding of vision along two essential axes: species differences and behavioral context. In freely moving marmoset monkeys, natural gaze behavior dynamically organizes retinal input by varying the timing and amplitude of gaze shifts, revealing V1 mechanisms that are obscured under head-fixed conditions where eye movements are constrained and decoupled from head motion. This perspective highlights vision as an active brain–behavior process, in which behavior structures sensory input and cortical circuits are optimized to process its resulting statistics. Although the present study establishes how natural gaze behavior organizes V1 responses, future work will be needed to determine how receptive-field computations and visual representations are further shaped under natural viewing conditions. Together, these insights move the field toward a naturalistic, behavior-centered framework for understanding real-world visual brain function and motivate future studies in freely moving primates to reveal how higher cortical areas integrate visual and non-visual signals during active behavior to structure the neural codes that give rise to our perception.

## Acknowledgements

This work was supported by NIH R01 NS118457 to C.T.M., NIH UF1 NS116377 and AFOSR 19RT0316 to C.T.M. and A.C.H., KIBM Postdoctoral Award and SFARI Fellows-to-Faculty Award to J.L.

## Author contribution

J.L., J.F.M., A.C.H., and C.T.M. designed research; J.L., and V.P.S. performed research; J.L. analyzed data; and J.L., J.F.M., A.C.H., and C.T.M. wrote the paper.

## Methods

### Subjects

Experiments in this work involved two adult common marmosets (*Callithrix jacchus*). Monkey S and G are both females and group housed. Monkey S had bilateral chronic implants in V1 and Monkey G had a chronic implant in left-hemisphere V1. Both subjects were at least 1.5 years old at time of implant. All surgeries and experiments were approved by the University of California San Diego (UCSD) Institutional Animal Care and Use Committee (IACUC) in accordance with National Institute of Health standards for care and use of laboratory animals.

### Neurophysiology

Neural activity was recorded with multi-electrode arrays (N-form, Modular Bionics, Berkeley, CA) chronically implanted in V1. The N-form array has 64 channels in total embedded in 16 shanks located in 4×4 grids evenly spaced 0.25 mm apart. Each shank has 4 iridium oxide electrodes located at 0.5, 0.375, 0.25, and 0 mm from the tip. During each recording session, a wireless Neurologger (SpikeLog-64, Deuteron Technologies) was hosted in a 3.5 cm (width) x 2.5 cm (height) x 1.2 cm (depth) protecting case and connected to the array to record the extracellular voltage at 32 kHz. Spike sorting was performed offline using Kilosort 2.0^50^ and manually curated using the graphic user interface Phy. A small number of neurons exhibiting abrupt waveform changes during manual curation were excluded from all analyses.

### Experiment design

In the freely moving paradigm, subjects were placed in a 2×3 m arena decorated with high-contrast visual stimuli of dots and gratings in 6 orientations (evenly-spaced 30-degree intervals) on the wall and floor. Subjects were allowed to freely move in the arena for 30 minutes. In the sessions with darkness, light was switched on and off every 5 minutes alternatively. When the light was switched off, all the remaining light sources, such as device signal lights, were well covered to ensure complete darkness. In the head-fixed paradigm, subjects were head-fixed viewing stimuli on the screen to map the V1 neurons’ receptive field and tuning property. For mapping receptive field, subjects freely viewed the stimuli of 120 randomly positioned black and white flashing dots generated at a frequency of 10 Hz for 5 minutes. For orientation and spatial frequency tuning, full-field drifting gratings with 12 orientations in 12 orientations (evenly-spaced 30-degree intervals) and 3 spatial frequencies (0.5, 1, 2 cycles/degree). Stimuli were displayed using MarmoV5. Subjects wear the head-mounted eye tracker and wireless Neurologger with continuous eye tracking and neural recording throughout the recording, including both head-fixed and freely moving conditions.

### Eye, head, body, and visual scene tracking

Eye and visual scene were recorded with the head-mounted eye tracker described in our previously published work^17^. Frame rates of the eye camera and visual scene camera are 90 fps and 60 fps, respectively. Pupil detection was performed using the previously developed algorithm UNet and custom-designed graphic user interface^17^. Eye calibration was performed offline by matching the pupil centroid to the visual scene images when subjects were head-fixed viewing marmoset faces shown on a screen. Head and body movements were recorded using the OptiTrack motion tracking system. Three infra-red reflective beads (12 mm diameter) were placed on the head-mounted eye tracker for the head rotation and movement, and a single bead was added to the backpack of eye tracker for body movement. Eight OptiTrack cameras were strategically placed in the arena and calibrated before every session (camera error under 0.1 mm) to obtain precise tracking of the markers at 120 fps. The tracking data were manually curated using the OptiTrack associated software Motive after the recording, where false markers were removed and gaps were filled with linear interpolation.

### Behavioral analysis

Eye and head position were first up-sampled to 200 Hz with linear interpolation. Then, gaze position was obtained from the sum of eye and head position. Gaze shifts and fixations were distinguished with a gaze velocity threshold of ±0.2 deg/ms. The threshold was chosen to well separate gaze shifts and fixations as eye and head are negatively correlated during fixations, while the negative correlation is broken during gaze shifts. To exclude the potential influence from other movement sources, time during locomotion was excluded in the analysis of the current manuscript. Locomotion was detected using the criteria: 1) body speed exceeded 3 cm/s after applying a 0.2 Hz low-pass filter; 2) the position of the head is lower than 18 cm; and 3) the duration of the period is larger than 3s. During the light condition, marmosets spend most of their time stationary, using locomotion primarily to approach a selected destination. In darkness, locomotion rarely occurred unless the animal was startled by an unexpected sound. Future studies that specifically dissociate locomotion-related signals from concurrent changes in visual input will be important for determining the role of locomotion in V1 activity in freely moving primates.

### PSTH analysis

Neurons with mean firing rate less than 1 Hz were excluded from the PSTH analysis. PSTHs were computed with 1 ms resolution in the analysis window of −200 to 400 ms relative to the event onset. A 10-fold jackknife procedure was applied to estimate PSTH mean and standard error. In each jackknife sample, PSTHs were smoothed using a Gaussian kernel with σ = 5 ms. Normalized PSTHs were computed by dividing the PSTH by its baseline and subtracting 1, where the baseline was defined as the mean within the pre-event window of −200 to −100 ms. Suppression and enhancement amplitudes were defined as the maximum dip and peak amplitudes of the normalized PSTH, respectively. Significance of suppression and enhancement was assessed by computing a z-score relative to the PSTH baseline; response with z-scores exceeding 2 were considered significant. Response latency was defined as the zero-crossing point between suppression and enhancement in the normalized PSTHs, as every neuron exhibited both suppression and enhancement regardless of statistical significance.

### Parametric models

We fit two parametric models, the saccade-locked model and two-stage model, to the normalized PSTHs. The parametric models assumed that suppression and enhancement components could each be approximated by a raised-cosine basis function. For the saccade-locked model, suppression and enhancement amplitude, width and latency were first extracted from the gaze-shift aligned normalized PSTH. For the two-stage model, suppression and enhancement parameters were extracted from the gaze-shift aligned and fixation aligned normalized PSTHs, respectively. These parameters were used to initialize a set of raised-cosine basis functions for model fitting:

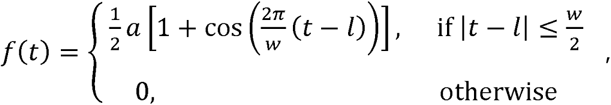

where *a*, *w*, and *l* represent amplitude, width and latency of suppression or enhancement, respectively. The basis functions were added to each gaze-movement event to produce a continuous neural response. In the saccade-locked model, both suppression and enhancement basis functions were added to gaze-shift onsets. In the two-stage model, suppression and enhancement were added to gaze-shift and fixation onsets, respectively. Then, predicted normalized PSTHs aligned to gaze-shift and fixation onsets were calculated from the continuous neural response. While widths and latencies were fixed parameters, suppression and enhancement amplitudes were optimized using constrained nonlinear multivariable minimization (fmincon, Matlab). The loss function was defined as the mean squared error between the

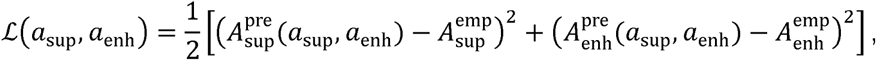

where *a* represents amplitude in basis functions; *A* represents amplitude in predicted and empirical PSTHs. For the saccade-locked model, *A_sup_* and *A_enh_* were both from gaze-shift aligned PSTH. For the two-stage model, *A_sup_* and *A_enh_* were from gaze-shift aligned and fixation aligned PSTH, respectively. The optimization stops when loss function fell below 10^−6^ or when the maximum of 20 iterations were reached. The performance of the parametric models was measured by the mean squared error between the predicted and empirical suppression and enhancement amplitudes in both gaze-shift aligned and fixation aligned PSTHs.

### Receptive field and preferred stimulus of V1 neurons

Receptive field and tuning of 251 V1 neurons from two monkeys were obtained in the head-fixed paradigm. Receptive field was mapped using reverse correlation, and a window of 5×5 degree^2^ was manually selected around the receptive field center. A weighted mean tuning in frequency domain, *F*_PREF_, was obtained to represent the preferred input for each V1 neuron. First, Fast Fourier Transform (FFT) was applied to stimulus conditions across different orientations and special frequencies. Next, the log-magnitude spectrum of FFT associated with each stimulus condition was weighted by the neuron’s corresponding firing rate, and these weighted components were summed across all conditions. The summed FFT log-magnitude spectrum was then normalized to the range of [0,1].

### Receptive field input in freely moving paradigm

In the freely moving paradigm, the visual scene during each fixation is extracted from the world-camera image captured by the head-mounted eye tracker. For each neuron, receptive field input was then cropped out from the visual scene in a 5×5 degree^2^ window centered on the receptive field position relative to the gaze center. Depending on the position of gaze center in the visual scene, the receptive field input might not fall within the image for the frames during a fixation, and thus not available. When multiple frames were present during a fixation, the second frame was chosen if available. If not, the third frame was selected when available, followed by the first frame. This ordering was chosen because the first frame was more likely to be influenced by residual head-motion artifacts near fixation onset. Fixations lacking any valid frame were excluded from the following analysis.

Next, receptive field input during fixations was mean-subtracted and added a raised-cosine filter before FFT was applied to obtain a frequency-domain representation. For each fixation, we calculated a predicted RF drive *D* by computing the normalized dot product of FFT log-magnitude spectrum of the neuron’s preferred RF input *F*_PREF_ and the current RF input *F_RF_*:

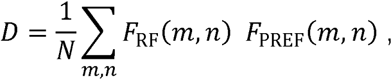

where *m*, *n*, and *N* denote the row, column, and total number of elements, respectively. For the analysis in Fig. 4, the top tertile *D* was defined as preferred input events, while the bottom tertile *D* was defined as non-preferred input events across all fixations throughout the recording session.

For the analysis in Fig. 5, the predicted RF drive was normalized to the range [0,1] for each neuron and recording session, after which data from all neurons and sessions were pooled. To exclude RF visual inputs with simply low- or high-contrast regardless of their spatial pattern, only values within the 10th–90th percentile of the normalized predicted RF drive distribution (0.41–0.77) were included in subsequent analyses. This range primarily reflects the alignment between the spatial pattern of RF visual input and the neuron’s tuning, rather than differences in stimulus contrast. For each neuron, the firing rate during fixation-associated enhancement was calculated within a 60-ms window centered on the enhancement latency, defined as the peak of enhancement in the fixation-aligned PSTH. A linear fit was then performed between the normalized predicted RF drive and the firing rate during fixation-associated enhancement. Neurons with a significant positive linear relationship (slope >0, p<0.05) were selected as representative examples for Fig. 5, whereas other neurons were shown in Fig. S4a-b.

RF input correlation was quantified as the Pearson correlation between the log-magnitude spectra of the fast Fourier transform (FFT) of receptive field (RF) inputs. Gaze shifts were classified as small (<3 deg), medium (3–13 deg), or large (>13 deg). These thresholds were chosen based on the RF size of the recorded V1 neurons (mostly <3 visual deg) and our previous finding that eye movements in freely moving marmosets are mostly within 13 deg^17^. Correlations for shuffled fixations were computed by randomly pairing fixations. Group comparisons were performed using paired t-tests with a significance threshold of p<0.05.

## References

1. Yarbus, A.L. (2013). Eye movements and vision (Springer).

2. Land, M., and Tatler, B. (2009). Looking and acting: vision and eye movements in natural behaviour (Oxford University Press).

3. Franchak, J.M. (2020). Visual exploratory behavior and its development. In Psychology of learning and motivation (Elsevier), pp. 59–94.

4. Ngo, V., Gorman, J.C., la Fuente, M.F., Souto, A., Schiel, N., and Miller, C.T. (2022). Active vision during prey capture in wild marmoset monkeys. Curr. Biol. 32, 3423–3428.

5. Parker, P.R.L., Martins, D.M., Leonard, E.S.P., Casey, N.M., Sharp, S.L., Abe, E.T.T., Smear, M.C., Yates, J.L., Mitchell, J.F., and Niell, C.M. (2023). A dynamic sequence of visual processing initiated by gaze shifts. Nat. Neurosci. 26, 2192–2202.

6. Skyberg, R.J., and Niell, C.M. (2024). Natural visual behavior and active sensing in the mouse. Curr. Opin. Neurobiol. 86, 102882.

7. Willmore, B.D.B., Prenger, R.J., and Gallant, J.L. (2010). Neural representation of natural images in visual area V2. J. Neurosci. 30, 2102–2114.

8. Nishimoto, S., and Gallant, J.L. (2011). A three-dimensional spatiotemporal receptive field model explains responses of area MT neurons to naturalistic movies. J. Neurosci. 31, 14551–14564.

9. Knöll, J., Pillow, J.W., and Huk, A.C. (2018). Lawful tracking of visual motion in humans, macaques, and marmosets in a naturalistic, continuous, and untrained behavioral context. Proc. Natl. Acad. Sci. 115, E10486--E10494.

10. Cadena, S.A., Denfield, G.H., Walker, E.Y., Gatys, L.A., Tolias, A.S., Bethge, M., and Ecker, A.S. (2019). Deep convolutional models improve predictions of macaque V1 responses to natural images. PLoS Comput. Biol. 15, e1006897.

11. Yates, J.L., Coop, S.H., Sarch, G.H., Wu, R.-J., Butts, D.A., Rucci, M., and Mitchell, J.F. (2023). Detailed characterization of neural selectivity in free viewing primates. Nat. Commun. 14, 3656.

12. Farzmahdi, A., Kohn, A., and Coen-Cagli, R. (2025). Relating natural image statistics to patterns of response covariability in macaque primary visual cortex. Nat. Commun. 16, 6757.

13. Angelaki, D.E., and Hess, B.J.M. (2005). Self-motion-induced eye movements: effects on visual acuity and navigation. Nat. Rev. Neurosci. 6, 966–976.

14. Matthis, J.S., Muller, K.S., Bonnen, K.L., and Hayhoe, M.M. (2022). Retinal optic flow during natural locomotion. PLOS Comput. Biol. 18, e1009575.

15. Muller, K.S., Matthis, J., Bonnen, K., Cormack, L.K., Huk, A.C., and Hayhoe, M. (2023). Retinal motion statistics during natural locomotion. Elife 12, e82410.

16. Hayhoe, M., Rothkopf, C., Goettker, A., and Powell, N. (2025). Natural visually guided behavior. Brain Res., 149899.

17. Singh, V.P., Li, J., Dawson, K., Mitchell, J.F., and Miller, C.T. (2025). Active vision in freely moving marmosets using head-mounted eye tracking. Proc. Natl. Acad. Sci. 122, e2412954122.

18. Mitchell, J.F., Reynolds, J.H., and Miller, C.T. (2014). Active vision in marmosets: a model system for visual neuroscience. J. Neurosci. 34, 1183–1194.

19. Solomon, S.G., and Rosa, M.G.P. (2014). A simpler primate brain: the visual system of the marmoset monkey. Front. Neural Circuits 8, 96.

20. Niell, C.M., and Stryker, M.P. (2010). Modulation of visual responses by behavioral state in mouse visual cortex. Neuron 65, 472–479.

21. Ayaz, A., Saleem, A.B., Schölvinck, M.L., and Carandini, M. (2013). Locomotion controls spatial integration in mouse visual cortex. Curr. Biol. 23, 890–894.

22. Saleem, A.B., Ayaz, A., Jeffery, K.J., Harris, K.D., and Carandini, M. (2013). Integration of visual motion and locomotion in mouse visual cortex. Nat. Neurosci. 16, 1864–1869.

23. Erisken, S., Vaiceliunaite, A., Jurjut, O., Fiorini, M., Katzner, S., and Busse, L. (2014). Effects of locomotion extend throughout the mouse early visual system. Curr. Biol. 24, 2899–2907.

24. Vinck, M., Batista-Brito, R., Knoblich, U., and Cardin, J.A. (2015). Arousal and locomotion make distinct contributions to cortical activity patterns and visual encoding. Neuron 86, 740–754.

25. Stringer, C., Pachitariu, M., Steinmetz, N., Reddy, C.B., Carandini, M., and Harris, K.D. (2019). Spontaneous behaviors drive multidimensional, brainwide activity. Science (80-. ). 364, eaav7893.

26. Musall, S., Kaufman, M.T., Juavinett, A.L., Gluf, S., and Churchland, A.K. (2019). Single-trial neural dynamics are dominated by richly varied movements. Nat. Neurosci. 22, 1677–1686.

27. Miura, S.K., and Scanziani, M. (2022). Distinguishing externally from saccade-induced motion in visual cortex. Nature 610, 135–142.

28. Duffy, F.H., and Burchfiel, J.L. (1975). Eye movement-related inhibition of primate visual neurons. Brain Res. 89, 121–132.

29. Kayama, Y., Riso, R.R., Bartlett, J.R., and Doty, R.W. (1979). Luxotonic responses of units in macaque striate cortex. J. Neurophysiol. 42, 1495–1517.

30. Kagan, I., Gur, M., and Snodderly, D.M. (2008). Saccades and drifts differentially modulate neuronal activity in V1: effects of retinal image motion, position, and extraretinal influences. J. Vis. 8, 19.

31. McFarland, J.M., Bondy, A.G., Saunders, R.C., Cumming, B.G., and Butts, D.A. (2015). Saccadic modulation of stimulus processing in primary visual cortex. Nat. Commun. 6, 8110.

32. Liska, J.P., Rowley, D.P., Nguyen, T.T.K., Muthmann, J.-O., Butts, D.A., Yates, J.L., and Huk, A.C. (2024). Running modulates primate and rodent visual cortex differently. 10.7554/elife.87736.2.

33. Talluri, B.C., Kang, I., Lazere, A., Quinn, K.R., Kaliss, N., Yates, J.L., Butts, D.A., and Nienborg, H. (2023). Activity in primate visual cortex is minimally driven by spontaneous movements. Nat. Neurosci. 26, 1953–1959.

34. Ibbotson, M., and Krekelberg, B. (2011). Visual perception and saccadic eye movements. Curr. Opin. Neurobiol. 21, 553–558.

35. Leopold, D.A., and Logothetis, N.K. (1998). Microsaccades differentially modulate neural activity in the striate and extrastriate visual cortex. Exp. Brain Res. 123, 341–345.

36. David, S. V, Hayden, B.Y., and Gallant, J.L. (2006). Spectral receptive field properties explain shape selectivity in area V4. J. Neurophysiol. 96, 3492–3505.

37. Diamond, M.R., Ross, J., and Morrone, M.C. (2000). Extraretinal control of saccadic suppression. J. Neurosci. 20, 3449–3455.

38. Ross, J., Morrone, M.C., Goldberg, M.E., and Burr, D.C. (2001). Changes in visual perception at the time of saccades. Trends Neurosci. 24, 113–121.

39. MacEvoy, S.P., Hanks, T.D., and Paradiso, M.A. (2008). Macaque V1 activity during natural vision: effects of natural scenes and saccades. J. Neurophysiol. 99, 460–472.

40. Ruiz, O., and Paradiso, M.A. (2012). Macaque V1 representations in natural and reduced visual contexts: Spatial and temporal properties and influence of saccadic eye movements. J. Neurophysiol. 108, 324–333.

41. Paradiso, M.A., Meshi, D., Pisarcik, J., and Levine, S. (2012). Eye movements reset visual perception. J. Vis. 12, 11.

42. Niemeyer, J.E., Akers-Campbell, S., Gregoire, A., and Paradiso, M.A. (2022). Perceptual enhancement and suppression correlate with V1 neural activity during active sensing. Curr. Biol. 32, 2654–2667.

43. Trotter, Y., and Celebrini, S. (1999). Gaze direction controls response gain in primary visual-cortex neurons. Nature 398, 239–242.

44. Morris, A.P., and Krekelberg, B. (2019). A stable visual world in primate primary visual cortex. Curr. Biol. 29, 1471–1480.

45. Guitchounts, G., Masís, J., Wolff, S.B.E., and Cox, D. (2020). Encoding of 3D head orienting movements in the primary visual cortex. Neuron 108, 512–525.

46. Denagamage, S., Morton, M.P., Hudson, N. V, Reynolds, J.H., Jadi, M.P., and Nandy, A.S. (2023). Laminar mechanisms of saccadic suppression in primate visual cortex. Cell Rep. 42.

47. Zanos, T.P., Mineault, P.J., Guitton, D., and Pack, C.C. (2016). Mechanisms of saccadic suppression in primate cortical area V4. J. Neurosci. 36, 9227–9239.

48. Miller, C.T., Gire, D., Hoke, K., Huk, A.C., Kelley, D., Leopold, D.A., Smear, M.C., Theunissen, F., Yartsev, M., and Niell, C.M. (2022). Natural behavior is the language of the brain. Curr. Biol. 32, R482--R493.

49. Laramée, M.-E., and Boire, D. (2015). Visual cortical areas of the mouse: comparison of parcellation and network structure with primates. Front. Neural Circuits 8, 149.

50. Pachitariu, M., Sridhar, S., Pennington, J., and Stringer, C. (2024). Spike sorting with Kilosort4. Nat. Methods, 1–8.

