## Supplemental for "Active gaze behavior organizes V1 activity in freely-moving marmosets"

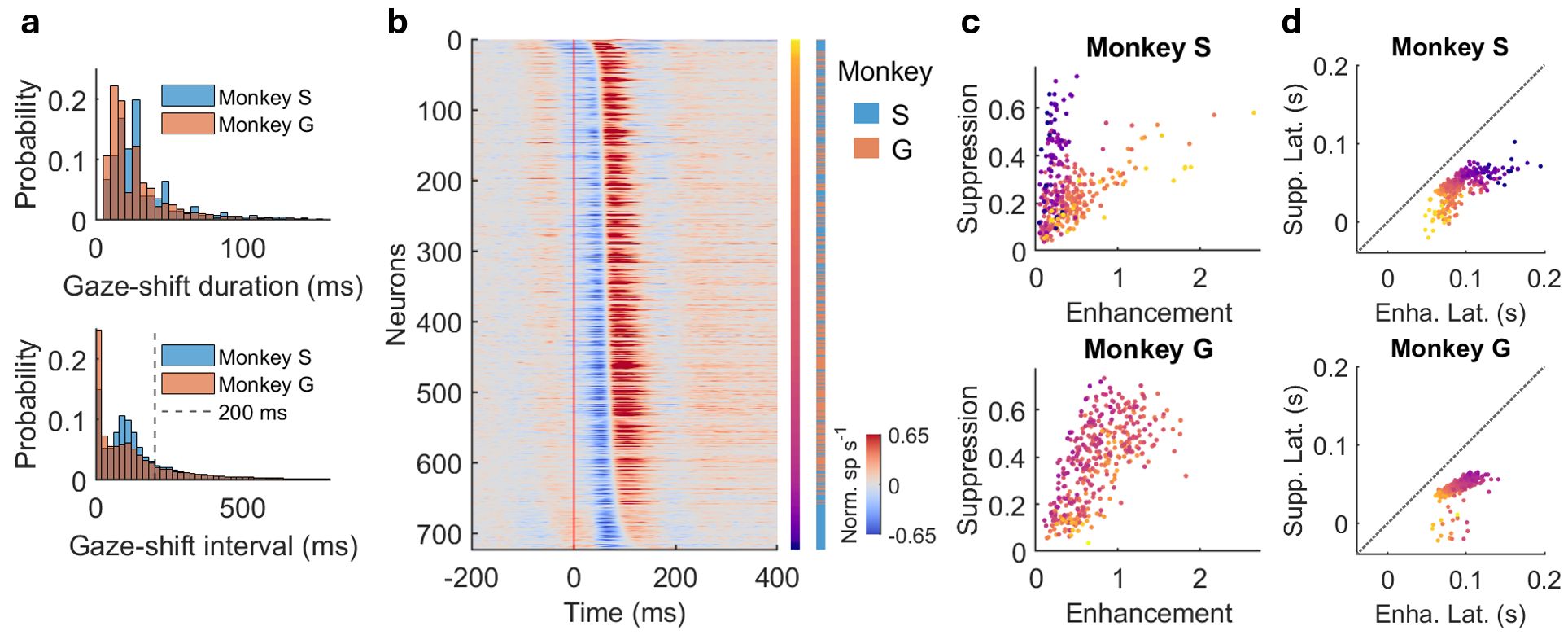

**Figure S1. Statistics of gaze-shift behavior and gaze-shift aligned neural responses across two monkeys.**

**a)** Histogram of gaze-shift duration (top) and interval (bottom) for the two monkeys. A large amount gaze-shift intervals fall within 200 ms, indicating frequent gaze shifts during active exploration of the 3D environment.

**b)** Normalized PSTHs aligned with gaze-shift onset (same as Fig. 1d) with identity of monkeys shown by the vertical bar.

**c)** Suppression amplitude versus enhancement amplitude (same as Fig. 1g) for each monkey. Each dot represents a neuron with color indicating latency as in **b**. The conclusion of larger suppression/enhancement ratio for later neurons than earlier neurons holds in each monkey.

**d)** Suppression latency versus enhancement latency for each monkey. Each dot represents a neuron with color indicating latency as in **b**. black doted lines indicate unity. Suppression and enhancement latencies are positively correlated across the population.

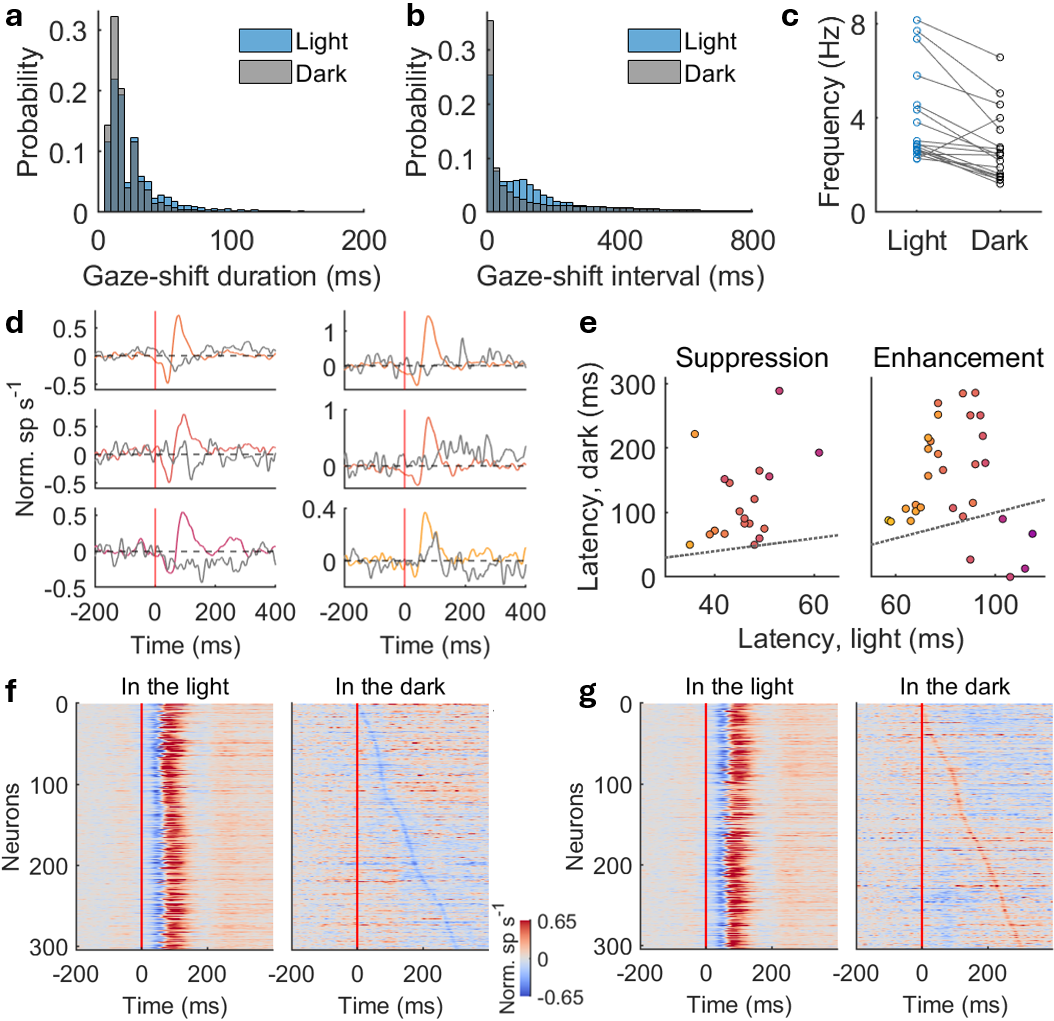

**Figure S2. Gaze behavior and neural responses in the light versus dark.**

**a)** Histogram of gaze-shift duration in the light and dark.

**b)** Histogram of gaze-shift interval in the light and dark.

**c)** Average frequency of gaze shifts in the light and dark for each session. Gaze shifts in the dark are generally less frequent than in the light but remain at a decent level.

**d)** Examples of neurons exhibiting significant suppression (left) and enhancement (right) in darkness. Colored and gray traces show responses in the light and dark conditions, respectively. Color indicates response latency as in Fig. 2e. Red vertical lines indicate gaze-shift onset. Responses in darkness exhibit substantially weaker and distinct temporal profiles compared with those observed in the light.

**e)** Response latency in the dark versus light for neurons exhibiting significant suppression (left) and enhancement (right) in the dark. Each dot represents a neuron with colors indicating response latency as in Fig. 2e. Black dotted lines indicate unity. Response latencies in darkness were not systematically correlated with those in the light.

**f)** Population responses sorted by suppression latency in the dark. Neurons were sorted by the latency of the minimum PSTH in the dark, regardless of statistical significance. Responses in the light are shown using the same neuron ordering.

**g)** Population responses sorted by enhancement latency in the dark. Neurons were sorted by the latency of the maximum PSTH in the dark, regardless of statistical significance. Responses in the light are shown using the same neuron ordering. The residual responses in the dark do not systematically follow the response dynamics observed in the light. Responses observed in the dark are substantially weaker and narrower than those observed in the light across the population.

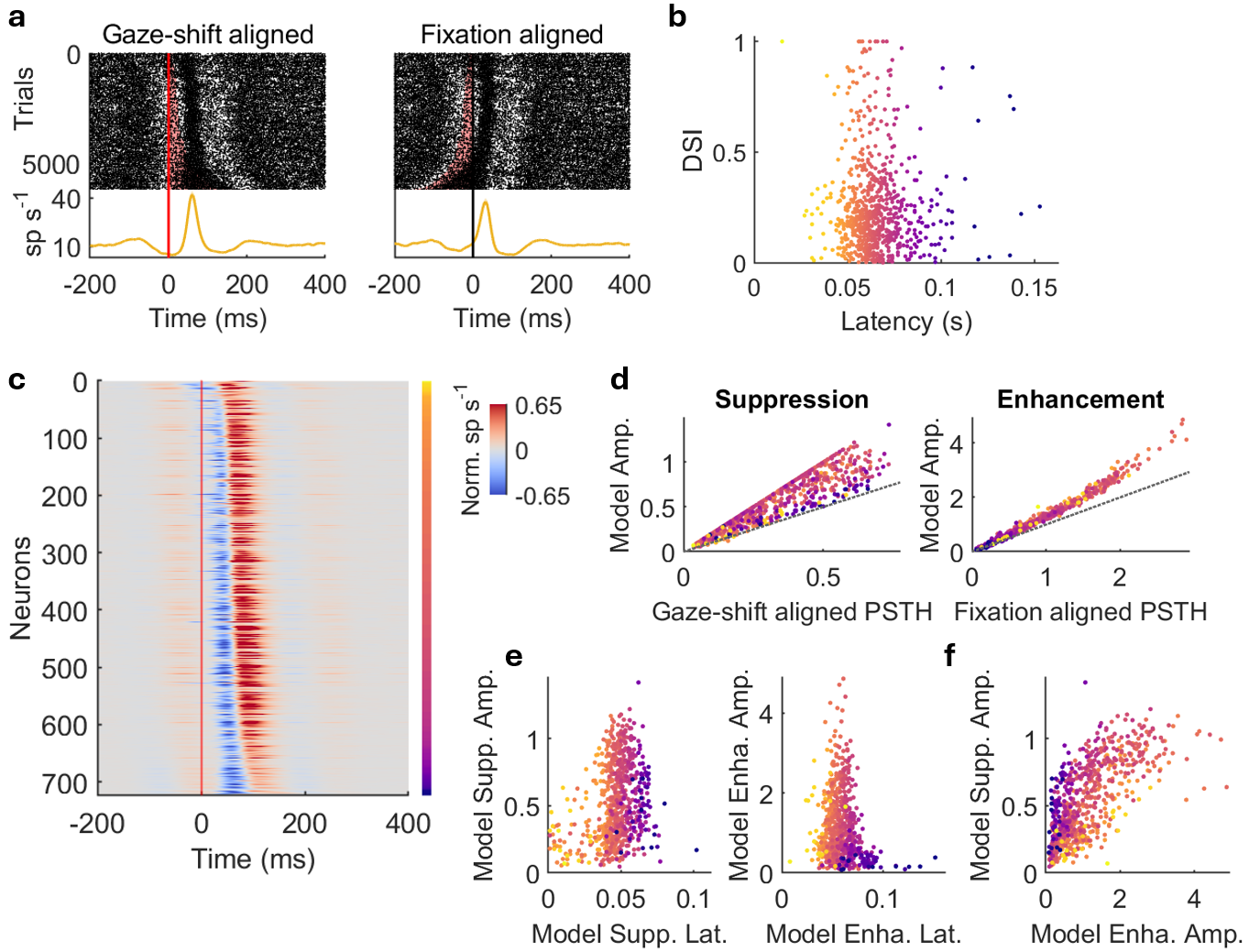

**Figure S3. Example early-latency neuron, model-obtained PSTHs aligned with gaze-shift onset, and model best-fit parameters in the V1 population.**

**a)** Raster plots and PSTHs aligned with gaze-shift onsets (left) and fixation onsets (right) for an example neuron with early latency. The enhancement locks in with gaze-shift onset instead of fixation onset in raster plots, and the enhancement amplitude is larger in the gaze-shift aligned PSTH than the fixation aligned PSTH.

**b)** Direction selectivity index (DSI) as a function of response latency. Each dot represents a neuron with color indicating response latency as in **c**. DSI is calculated as (R_pref_-R_null_)/(R_pref_+R_null_), where Rpref and are the spike rate during preferred stimuli in drifting gratings and Rnull is the spike rate during the opposite direction to preferred stimuli. DSI was not significantly correlated with response latency across the population.

**c)** Normalized PSTHs aligned with gaze-shift onset obtained from the two-stage model. The neurons are sorted by response latencies same as Fig. 1d. The two-stage model, which incorporates the actual sequence of gaze events, recapitulates not only the main response but also the weak suppression and enhancement components flanking the main response.

**d)** Left: best-fit suppression amplitude from the model as a function of the suppression amplitude in gaze-shift aligned PSTH. Right: best-fit enhancement amplitude from the model as a function of the enhancement amplitude in fixation aligned PSTH. Each dot represents a neuron with color indicating response latency as in **c**. Black dotted lines indicate unity. Amplitudes from the model are larger than the measured PSTH amplitudes, indicating that temporal overlap between suppression and enhancement reduces their apparent amplitudes in the PSTHs.

**e)** Left: best-fit suppression amplitude from the model as a function of suppression latency used in the model. Right: best-fit enhancement amplitude from the model as a function of enhancement latency used in the model. Each dot represents a neuron with color indicating response latency as in **c**. Best-fit suppression amplitude was positively correlated with suppression latency, whereas best-fit enhancement amplitude was negatively correlated with enhancement latency.

**f)** Best-fit suppression amplitude versus enhancement amplitude from the model. Each dot represents a neuron with color indicating latency as in **c**. The amplitudes exhibited the same relationship as the corresponding amplitudes measured from the PSTHs in Fig. 1g.

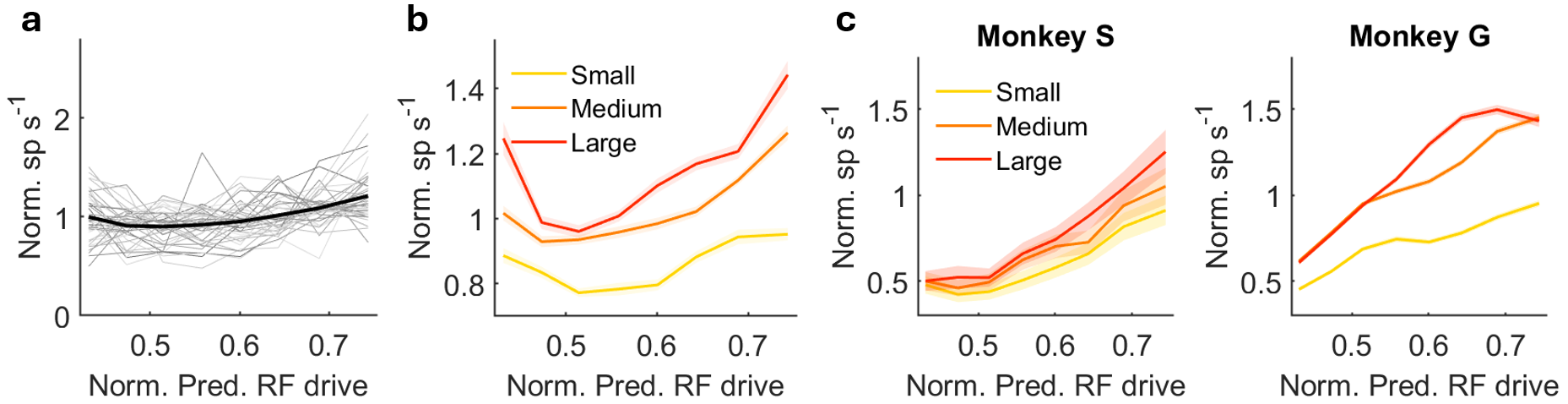

**Figure S4. Other neurons not included in the main analysis, and validation of the effect across individual monkeys.**

**a)** Normalized firing rate as a function of normalized predicted RF drive for other neurons that are not included in Fig. 5c (non-positive linear relationship or non-significant positive linear relationship). Gray lines show individual neurons; black line represents the population mean. Firing rates are normalized to each neuron's mean response across normalized predicted RF drive bins.

**b)** Population average of normalized firing rate following fixation as a function of normalized predicted RF drive for other neurons as in **a**, grouped by the amplitude of the preceding gaze shift. Firing rates are normalized to each neuron's mean response across preceding gaze-shift amplitudes and normalized predicted RF drive bins. Shades represent standard error.

**c)** Population average of normalized firing rate following fixation as a function of normalized predicted RF drive as in Fig. 5g for each monkey. The conclusion of firing rates increase with larger amplitude of the preceding gaze shift across predicted RF drive values holds in each monkey.

**Table S1. Key resources table.** **All original code, behavioral and neural data have been deposited at Dryad and are publicly available as of the date of publication.**

| REAGENT or RESOURCE | SOURCE | IDENTIFIER |
| --- | --- | --- |
| Deposited data | | |
| Behavioral and neural data | Dryad | DOI: available upon acceptance |
| Experimental models: Organisms/strains | | |
| Marmoset | University of California San Diego | N/A |
| Software and algorithms | | |
| Matlab 2021a | MathWorks | https://www.mathworks.com/products/new_products/release2021a.html |
| Kilosort 2.0 | Kilosort | https://github.com/jamesjun/Kilosort2 |
| Phy | Phy | https://phy.readthedocs.io/en/latest/ |
| Motive | OptiTrack | https://optitrack.com/software/motive/ |
| MarmoV5 | GitHub | https://github.com/jcbyts/MarmoV5 |
| UNet for pupil detection | GitHub | https://github.com/Vickey17/UNET_implementation_V2 |
| Code | Dryad | DOI: available upon acceptance |
